# Speed breeding: a powerful tool to accelerate crop research and breeding

**DOI:** 10.1101/161182

**Authors:** Amy Watson, Sreya Ghosh, Matthew J. Williams, William S. Cuddy, James Simmonds, María-Dolores Rey, M. Asyraf Md Hatta, Alison Hinchliffe, Andrew Steed, Daniel Reynolds, Nikolai Adamski, Andy Breakspear, Andrey Korolev, Tracey Rayner, Laura E. Dixon, Adnan Riaz, William Martin, Merrill Ryan, David Edwards, Jacqueline Batley, Harsh Raman, Christian Rogers, Claire Domoney, Graham Moore, Wendy Harwood, Paul Nicholson, Mark J. Dieters, Ian H. DeLacy, Ji Zhou, Cristobal Uauy, Scott A. Boden, Robert F. Park, Brande B. H. Wulff, Lee T. Hickey

**Author notes:** The contribution of these authors should be considered equal (Amy Watson, Sreya Ghosh).

## Abstract

The growing human population and a changing environment have raised significant concern for global food security, with the current improvement rate of several important crops inadequate to meet future demand [1]. This slow improvement rate is attributed partly to the long generation times of crop plants. Here we present a method called ‘speed breeding’, which greatly shortens generation time and accelerates breeding and research programs. Speed breeding can be used to achieve up to 6 generations per year for spring wheat (*Triticum aestivum*), durum wheat (*T. durum*), barley (*Hordeum vulgare*), chickpea (*Cicer arietinum*), and pea (*Pisum sativum*) and 4 generations for canola (*Brassica napus*), instead of 2-3 under normal glasshouse conditions. We demonstrate that speed breeding in fully-enclosed controlled-environment growth chambers can accelerate plant development for research purposes, including phenotyping of adult plant traits, mutant studies, and transformation. The use of supplemental lighting in a glasshouse environment allows rapid generation cycling through single seed descent and potential for adaptation to larger-scale crop improvement programs. Cost-saving through LED supplemental lighting is also outlined. We envisage great potential for integrating speed breeding with other modern crop breeding technologies, including high-throughput genotyping, genome editing, and genomic selection, accelerating the rate of crop improvement.

---

For most crop plants, the breeding of new, advanced cultivars, takes several years. Following crossing of selected parent lines, 4-6 generations of inbreeding are typically required to develop genetically stable lines for evaluation of agronomic traits and yield. This is particularly time-consuming for field-grown crops that are often limited to only 1-2 generations per year. Here, we present flexible protocols for “speed breeding” that use prolonged photoperiods to accelerate the developmental rate of plants [2], thereby reducing generation time. We highlight the opportunity presented by speed breeding and detail protocols to inspire widespread adoption as a state-of-the-art breeding and research tool.

To evaluate speed breeding as a method to accelerate applied and basic research on cereal species, standard genotypes of spring bread wheat (*T. aestivum*), durum wheat (*T. durum*), barley (*H. vulgare*) and the model grass *Brachypodium distachyon* were grown in a controlled environment room with extended photoperiod (22 hours light/2 hours dark) (Fig. 1; Methods: Speed breeding I; Supplementary Table 1). A light/dark period was chosen over a continuous photoperiod to support functional expression of circadian clock genes [3]. Growth was compared with that of plants in glasshouses with no supplementary light or heating during the spring and early summer of 2016 (Norwich, UK). Plants grown under speed breeding progressed to anthesis (flowering) in approximately half the time of those from glasshouse conditions. Depending on the cultivar or accession, anthesis was reached in 37 to 39 days (wheat - with the exception of Chinese Spring), and 37 to 38 days (barley), while it took 26 days to reach heading in *B. distachyon* (Fig. 2a,b,c,d; Supplementary Tables 2-4). Concurrently, the corresponding glasshouse plants reached the early stem-elongation growth stage or 3-leaf stage respectively. Wheat seed counts per spike decreased, although not always significantly, in the speed breeding chamber compared to the glasshouse with no supplementary light (Supplementary Table 5) and both wheat and barley plants produced a healthy number of spikes per plant, despite the rapid growth (Supplementary Table 6). Viability of mature seeds was unaffected by speed breeding with similar seed germination rates observed for all species (Supplementary Table 7). Moreover, crosses made between wheat cultivars under speed breeding conditions produced viable seed, including crosses between tetraploid and hexaploid wheat (Supplementary Table 8). These conditions were also used to successfully reduce the generation time of the model legume *Medicago truncatula* and the rapid cycling pea (*Pisum sativum*) variety JI 2822 [4] (Supplementary Tables 9, 10; Supplementary Fig. 1a,b).

**Figure 1.**
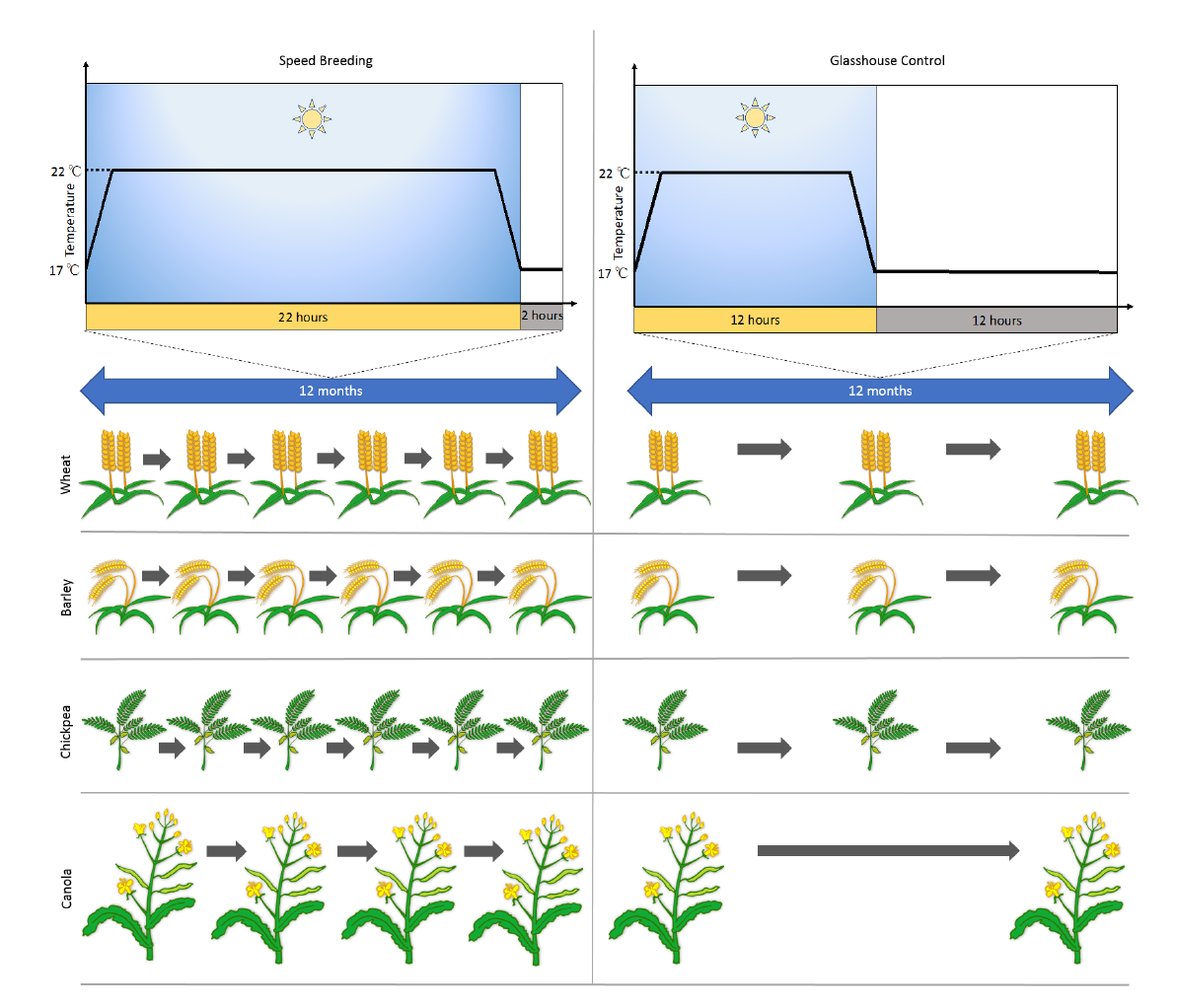
Speed breeding accelerates generation time of some major crop plants for research and breeding. Compared to a glasshouse with a natural photoperiod, where only 2-3 generations of wheat, barley, chickpea and canola can be achieved per year (right), speed breeding enables 4-6 generations of these crops to be grown in a year (left).

**Figure 2.**
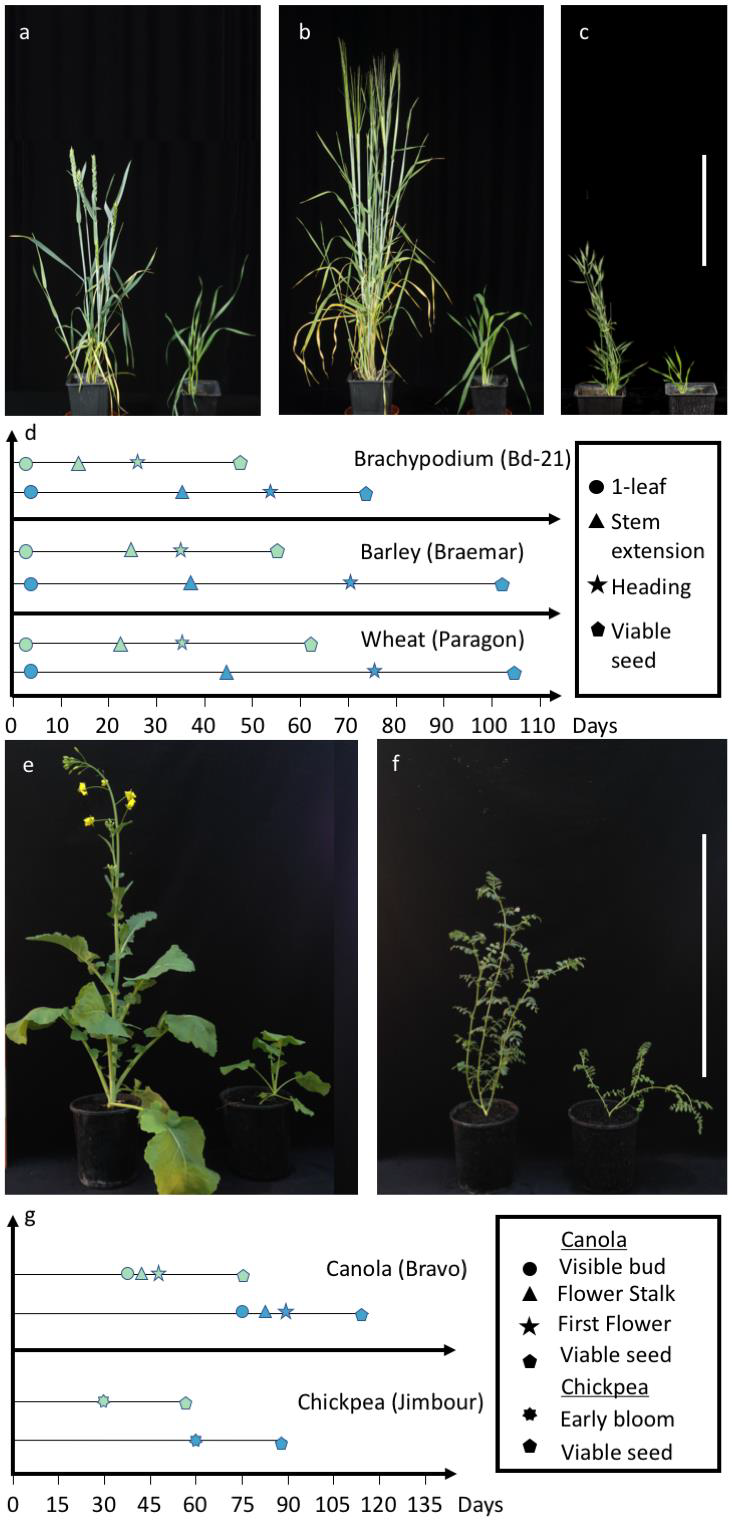
Accelerated plant growth and development under speed breeding (left) compared to control (right) conditions. **a**, Wheat (*T. aestivum cv*. Cadenza) at 38 days post sowing. **b**, Barley (*H. vulgare cv*. Braemar) at 41 days post sowing. **c**, Brachypodium (*B. distachyon* accession Bd21) at 36 days post sowing. Scale bar = 20 cm. **d**, Representative graph depicting the development stages of wheat, barley and brachypodium (x-axis in days) under speed breeding (top graph for each crop) conditions in a controlled environment chamber compared to control conditions (bottom graph for each crop) in a glasshouse in UK summer with no supplementary light or heat. **e**, Canola (*B. napus cv*. Bravo), pictured at 50 days post sowing. Scale bar = 50 cm. **f**, Chickpea (*C. arietinum cv*. Jimbour), pictured at 35 days post sowing. **g**, Representative graph depicting the development stages of canola and chickpea (x-axis in days) under speed breeding conditions in a supplemented glasshouse (top graph for each crop) compared to control conditions (bottom graph for each crop) in a glasshouse in Queensland, Australia, with no supplementary light or heating.

In an alternative, yet similar protocol for rapid generation cycling, we evaluated a collection of spring wheat, barley, canola and chickpea varieties in Queensland, Australia, in a temperature-controlled glasshouse fitted with high pressure sodium lamps to extend the photoperiod to a day-length of 22 hours. A control treatment in a glasshouse used a natural 12-hour control photoperiod. Both used the same temperature regime (22/17°C; Fig. 1; Methods: Speed breeding II; Supplementary Table 1). Time to anthesis was significantly reduced for all crop species relative to the 12-hour day-neutral photoperiod conditions, where the average reduction was, depending on genotype, 22 ± 2 days (wheat), 64 ± 8 days (barley), 73 ± 9 days (canola) and 33 ± 2 days (chickpea) (Fig. 2e-g; Supplementary Tables 11-14). Analysis of growth stage progression revealed normal, though accelerated, development for all species (Fig. 2e-g; Supplementary Tables 15-18) compared to the day-neutral conditions. Wheat plants produced significantly more spikes than those in day-neutral conditions and grain number was unaffected by the rapid development in both wheat and barley (Supplementary Tables 19, 20). Notably, time to anthesis was more uniform within each species under speed breeding conditions (Supplementary Tables 11-14 SD), an important feature, as synchronous flowering across genotypes is desirable for crossing. Additionally, wheat seed was harvested before maturity: 14 days post-anthesis in speed breeding conditions and following a 4-day cold treatment seed viability was high (Supplementary Table 21) indicating generation time can be further reduced by harvesting premature seed without the need for labour intensive embryo rescue [5]. Seed viability of all other species under Speed breeding II conditions were either unaffected or improved compared with day-neutral conditions (Supplementary Tables 22-24). Since temperature greatly influences the rate of plant development [6], generation time may be further accelerated by elevating temperature. This may however induce stress and affect plant performance.

Seed production (g/plant) of canola and chickpea was similar between speed breeding and day-neutral conditions (Supplementary Tables 25, 26). Considering the time taken to produce viable seed under speed breeding in the glasshouse, wheat, barley, canola and chickpea could produce 5.7, 5.4, 3.8 and 4.5 generations per year, respectively (Supplementary Table 27). These results highlight the opportunity to apply speed breeding across several important crop species without jeopardising the production of subsequent generations. The application of speed breeding conditions in a glasshouse fitted with supplementary lighting exemplifies the flexibility of the approach and may be preferred over growth chambers if rapid generation advance is to be applied to large populations, such as in breeding programmes.

Single seed descent (SSD) is commonly used in breeding programs and research to facilitate development of homozygous lines following a cross [7]. This process only requires one seed per plant to advance each generation. To investigate the ability of speed breeding to accelerate SSD, where plants are typically grown at high density, wheat and barley genotypes were grown in 100-cell trays under both speed breeding and 12-hour day-neutral photoperiod conditions in the glasshouse. This equated to a density of approximately 900 plants per m^2^. Generation time was shorter than for plants grown at lower density in the previous speed breeding experiments (Supplementary Tables 28, 29). This was likely caused by stress or plant competition as a result of the higher density, which is known to hasten flowering [8]. In most cases, each plant produced a single viable tiller containing on average 19.0 ± 1.3 and 18.6 ± 1.8 seeds for wheat and barley, respectively. Seed viability was 80% for wheat at two weeks post-anthesis and 100% for both wheat and barley at four weeks post anthesis (Supplementary Tables 28, 29). Therefore, integrating speed breeding and SSD techniques can effectively accelerate the generation of inbred lines for research and plant breeding programs.

In an additional protocol (Supplementary Table 1; Speed breeding III), we successfully implemented a low cost speed breeding growth room design, lit exclusively by LEDs (light-emitting diode) to reduce the operational cost of lighting and cooling, which permits 4-5 generations a year, depending on genotype and crossing plans (Supplementary Fig. 2). These results highlight the flexibility of tailoring the speed breeding ‘recipe’ to suit the local purpose and resources.

Besides exploring the various ways in which speed breeding can be used to accelerate generation time, we also evaluated the ability to phenotype some key adult plant phenotypes of wheat and barley. We observed that phenotypes associated with the EMS-induced mutation of the awn suppressor *B1* locus [9] and the Green Revolution *Reduced height* (*Rht*) genes in wheat [10] could be accurately recapitulated in the controlled environment room conditions (Fig. 3a,b). We also evaluated the effects that speed breeding might have on disease by inoculating resistant (Sumai 3) and susceptible (Timstein) wheat spikes with *Fusarium graminearum*, the causal agent of fusarium head blight (FHB). Consistent with expectations, we found clear signs of FHB progression in the susceptible cultivar, and little to no disease progression in the resistant cultivar (Fig. 3c; Supplementary Table 30). Previously, it has been shown that adult plant resistance (APR) to wheat leaf rust (caused by *Puccinia triticina* f. sp. *tritici*) and wheat stripe rust (*Puccinia striiformis* f. sp. *tritici*) can also be scored accurately under speed breeding conditions [11, 12]. We also evaluated the effect of loss-of-function of *FLOWERING LOCUS T-B1* in the F6 RIL of Paragon x W352 under speed breeding conditions. We observed the expected late flowering phenotype in the RIL, albeit within fewer days and with fewer leaves produced (Fig. 3d; Supplementary Table 31). To study the effect of speed breeding on cuticular β-diketone wax production in barley, we grew wild type and *Eceriferum* mutants [13]. The mutant leaf sheaths exhibited a clear, decreased glaucous appearance in the flag leaf stage, compared to the control (Fig. 3e). We also studied polyploid wheat chromosome pairing at metaphase I in meiosis in the presence and absence of the *PAIRING HOMOEOLOGOUS 1* (*Ph1*) locus [14]. We grew wheat carrying *Ph1* and wheat-rye hybrids either carrying or lacking *Ph1* in speed breeding and control conditions, and observed no major differences in chromosome pairing and recombination in meiocytes at metaphase I (Supplementary Table 32). The chromosome behavior suggests that both wheat and wheat-rye hybrids are cytologically stable under speed breeding conditions (Supplementary Fig. 3). In summary, all the above adult plant phenotypes could be recapitulated accurately and much faster than in the corresponding glasshouse conditions.

**Figure 3.**
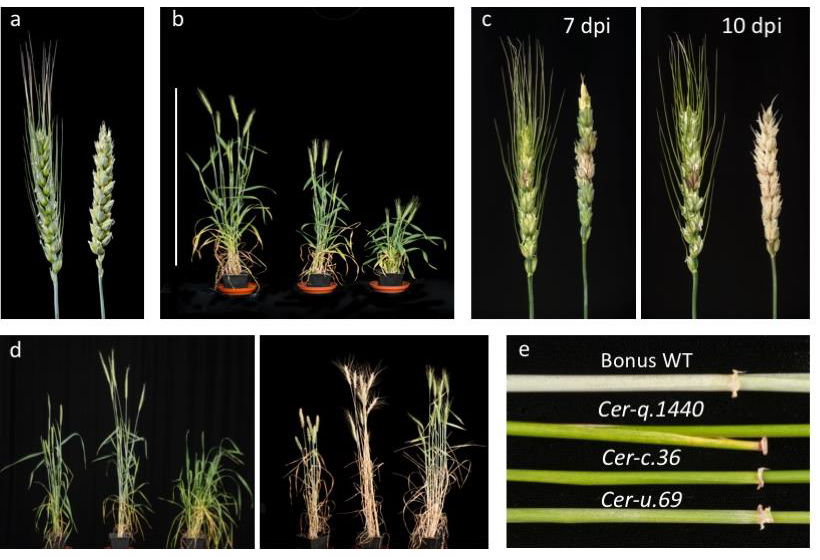
Adult plant phenotypes in wheat and barley under speed breeding conditions. **a**, Loss of function of the awn suppressor *B1* locus in *T. aestivum cv*. Paragon (mutant on left, wild type on right). **b**, Green Revolution *Reduced height* (*Rht*) dwarfing genes under speed breeding conditions. From left to right, *T. aestivum cv*. Maringá wild type, Maringá *Rht-1*, Maringá *Rht-3*. Scale bar = 100 cm. **c**, Fusarium Head Blight progression in spikes of the resistant *T. aestivum cv*. Sumai 3 (right) and the susceptible *cv*. Timstein (left) at 7 and 10 days post inoculation (dpi). **d**, Two instances depicting the flowering time of the F_6_ hybrid (right in both panels) derived from *T. aestivum cv*. Paragon (left in both panels) and *T. aestivum* landrace W352 (centre in both panels). In the left panel, the parents flower while the F_6_ is just beginning to boot at 44 days; in the right panel, the F_6_ hybrid is shown with developed spikes at 67 days when the parents have moved into senescence. **e**, Reduced glaucousness of *Eceriferum (Cer)* mutants in *H. vulgare cv*. Bonus.

We further investigated the potential for using speed breeding in conjunction with genetic transformation. Seeds of barley (*cv*. Golden Promise) were sown and immature embryos harvested and transformed (Supplementary Fig. 4). The transformation efficiency of speed grown and normal plants was comparable (Supplementary Table 33). Moreover, we observed that transformed barley explants could be grown under speed breeding conditions and viable seed obtained >6 weeks earlier than in standard control conditions (Supplementary Table 34; Supplementary Fig. 5).

We investigated the ability to phenotype canola, grown under glasshouse speed breeding, conditions for pod shattering resistance, a significant factor causing yield loss in the field. Using an established method which measures the energy absorbed by each pod when shattered by a pendulum [15] (Supplementary Fig. 6) we tested five canola cultivars considered susceptible to shattering. All displayed varying degrees of pod strength comparable to measures obtained from field-grown pods (Supplementary Table 35). Due to the lack of shatter resistance in cultivars, new alleles from related Brassica species could be transfered to canola [16]. Our results suggest that speed breeding could accelerate this objective.

Speed breeding is likely to reduce generation time for other crop species, such as sunflower (*Helianthus annuus*), pepper (*Capsicum annuum*), and radish (*Raphanus sativus*), which have been shown to respond well to extended photoperiod [2]. Speed breeding methods have already been successfully applied to accelerate breeding objectives for amaranth (*Amaranthus* spp.) [17] and peanut (*Arachis hypogaea*) [18]. For species that require short days to trigger the reproductive phase, such as rice (*Oryza sativa*) and maize (*Zea mays*), the speed breeding technique could be used to promote rapid vegetative growth prior to reducing the photoperiod.

Recent advances in genomic tools and resources [19-23] and the decreasing cost of sequencing have enabled plant researchers to shift their focus from model to crop plants. Despite such advances, the slow generation times of many crop plants continue to impose a high entry barrier. We envisage that combining these tools and resources with speed breeding will provide a strong incentive for more plant scientists to perform research on crop plants directly, thus further accelerating crop improvement research. In a breeding context, rapid generation advance to homozygosity following crossing will facilitate genetic gain for key traits and allow more rapid production of improved cultivars by breeding programs.

## Methods

### Speed breeding I - Controlled environment chamber speed breeding conditions

A Conviron BDW chamber (Conviron, Canada) was programmed to run a 22-hour photoperiod, with a temperature of 22 °C during the photoperiod, and 17 °C during the 2-hour dark period. Light and temperature were set to ramp up and down for 1 hour 30 minutes to mimic natural dawn and dusk conditions (as illustrated in Fig. 1). Humidity was set to 70%. Lighting was supplied by a mixture of white LED bars (Valoya; 6 units per 3.67 m^2^), far red LED lamps (Valoya; 12 units per 3.67 m^2^) and metal HQI lamps (Valoya; 32 units per 3.67 m^2^). Light intensity was adjusted to 360-380 µmol m^-2^ s^-1^ (highest value after ramping) at bench height, where the pots were kept, and 490-500 µmol m^-2^ s^-1^ (highest value after ramping) at adult plant height (with reference to wheat *cv.* Paragon). Light quality (spectral composition) is shown in Supplementary Fig. 7.

A timelapse video of *Triticum aestivum cv.* Paragon under speed breeding condition I (“Speed breeding”) and control glasshouse conditions (“control conditions”) in UK Summer with no supplementary light in the spring and summer of 2017 is given in Supplementary Media File 1 and the accompanying plant height growth curve in Supplementary Fig. 8.

### Sowing, harvesting, and germination rate evaluation for speed breeding in a controlled environment room chamber

Wheat, barley and *Brachypodium* seeds were imbibed on filter paper wetted with sterile water containing 0.5 ppm gibberellic acid, stratified at 4 °C for 3 days and then germinated at room temperature for 2 days before sowing in 1.33 L pots containing 900 mL of soil (one seed per pot). The soil used was the John Innes Centre (JIC) cereal mix, the composition of which is outlined in Supplementary Table 36. Seeds were harvested ~2.5-3 weeks post anthesis. The plants were not watered in the last week leading up to harvest in order to accelerate the grain ripening. The harvested spikes were kept in paper bags and dried at 28-30 °C for 3-5 days to reduce the moisture content. The grain was threshed and stored at room temperature. To measure germination rates, seeds were placed in Petri dishes lined with filter paper, and wetted with sterile water. The seeds were stratified for 4 °C for 3 days, and then placed at room temperature, with periodic rehydration of the filter paper lining to maintain moisture levels. Seeds were observed for germination at daily intervals for 5 days.

Pea seeds of the John Innes accession JI 2822 were scarified and sown into pots of approximately 260 ml volume with JIC Cereal Compost Mix (Supplementary Table 36). Pots were placed in a glasshouse with no supplementary light and a minimum temperature of 12 °C or the Conviron chamber under speed breeding conditions. The number of days after sowing until the first flower opened, flowering node number, number of pods, plant height and seed number, were recorded (Supplementary Table 10). At six weeks and nine weeks, respectively, the plants in the cabinet and glasshouse were no longer watered. Mature, dry seed were chipped and placed on moist filter paper for 5 days at room temperature in the dark to estimate percentage germination. The number of seeds where the radicle and shoot had not emerged were counted as not germinated.

Wild-type *M. truncatula* (genotype Jemalong A17) seeds were scarified with glasspaper, surface-sterilized for 3 minutes in 10% sodium hypochlorite and washed five times with distilled water. Seeds were left to imbibe for 2 hours before being transferred to agar plates for 72 hours at 4 °C. Seedlings were sown in JIC Cereal Compost Mix in 24-cell trays (each cell containing 8 0 mL of soil) and transferred to 1 L pots after 14 days. The number of days after sowing for the first flower to appear was recorded. Watering stopped following maturation of ~90% of the pods. Pods were harvested 14 days later and the resulting seeds subjected to germination tests similar to pea.

### Speed breeding II - Glasshouse speed breeding conditions

A temperature-controlled glasshouse fitted with high pressure sodium vapour lamps (Philips SON-T 400W E E40 in Sylvan High Bay housing with glass diffuser) was programmed to a 17/22 °C temperature regime with a 12-hour turnover and 22-hour photoperiod (Supplementary Fig. 9). The 2-hour period without lamps operating and 17 °C cycle occurred during the night. Light intensity was 440-650 µmol m^-2^ s^-1^ at adult plant height (approximately 45 cm above bench height). Light and temperature changes did not include a ramping up/down procedure. Light quality (spectral composition) is shown in Supplementary Fig. 10.

### Sowing, harvesting, and germination rate evaluation for speed breeding in a light-supplemented glasshouse

Wheat, barley, canola and chickpea seeds were imbibed by placing on water-moistened filter paper in agar plates overnight at room temperature. The imbibed seeds were chilled at 4 °C for 4 days to break any dormancy and support uniform germination. Seeds were then left at room temperature for 5 days. All seedlings were sown singly into 1.4 L pots. Soil media consisted of the CGS20 compost mix (Supplementary Table 37) with the addition of Scotts Osmocote^R^ Plus trace elements (2 g/L soil) and a final pH of 5.5-6.5. Pots were placed in pot racks in complete randomized block designs and hand-watered daily. Plants were supplemented with a weekly foliar spray of calcium nitrate diluted in filtered water (1 g/L) to mitigate any calcium deficiency, which is common with such rapid growth [24].

Anthesis date was recorded for wheat, canola and chickpea while awn peep was recorded for barley. One spike was harvested from wheat and barley at 2 and 4 weeks post-anthesis. Three pods per plant of canola and chickpea were harvested 6 weeks post-anthesis. All harvested seeds were placed in paper bags and immediately dried in an air-forced oven for 5 days at 35 °C, threshed by hand, then weighed. Germination tests were carried out as per the pre-sowing treatments described above. Percentage germination was calculated following 5 days at room temperature. At approximately 4-6 weeks following anthesis, water supply was reduced to every second day for one week, then twice per week for one week, followed by complete withholding of water. When plants were fully senesced, all spikes or pods were harvested, counted, dried, threshed and weighed to determine yield. Grain number per spike of wheat and barley plants was counted following threshing of each individual spike.

One cultivar each of wheat (*cv*. Westonia) and barley (*cv*. Commander) was sown in a 100-cell tray (length, 350 mm; width, 290 mm; height 45 mm; each cell approximately 18 ml) to demonstrate the potential of high density planting. Three spikes were harvested at 2 and 4 weeks post-anthesis and percentage germination determined, as above. The watering regime was also as above. Following complete senescence, all grain was harvested, dried, threshed and weighed to determine yield.

As a control, an identical protocol was carried out on the same species/cultivars in a similar glasshouse maintained at the same temperature settings but with no supplemental lighting.

### Speed breeding III - Homemade growth room design for low cost speed breeding

A room of 3 m x 3 m x 3 m with insulated sandwich panelling fitted with 7 LB-8 LED light boxes (1 light box per 0.65 m^2^) from Grow Candy (www.growcandy.com) and a 1.5 hp inverter split system domestic air conditioner was set up as a low cost alternative to the Conviron BDW chamber. The light spectrum details are outlined in Supplementary Fig. 11. The light quantity of photosynthetically active radiation (PAR) at bench height ranged from 210-260 µmol m^-2^ s^-1^ and at 50 cm above the pot from 340-590 µmol m^-2^ s^-1^. The lights were situated at a height of 140 cm above the bench. The room can accommodate 90 pots of 8” diameter and 5 L volume. Automatic watering was achieved with the Hunter 10 Station Irrigation Controller, with one solenoid per room and a 13 mm main line with spike drippers (one per 8” pot). The humidity conditions were ambient.

The lighting was set to run a 12-hour photoperiod (12 hours darkness) for 4 weeks and then increased to an 18-hour photoperiod (6 hours darkness). The lights did not ramp up and down during the switch between light and dark periods in the 24-hour cycle. The air-conditioner was set to run at 18 °C in darkness and 21 °C when the LED lights were on, with fluctuation being no more than ± 1 °C.

This system was used for speed breeding of wheat (*T. aestivum cvs*. Morocco and AvocetR), barley (*H. vulgare cvs.* Gus and Baudin), oat (*Avena sativa cv*. Swan), and triticale (*Triticosecale cvs*. Jackie and Coorong).

### Adult phenotyping protocols

#### Canola pod shattering resistance phenotyping

Five canola cultivars were selected for evaluation (*cvs.* Skipton, CB Argyle, ATR Cobbler, ATR Beacon and Bravo TT), where 10 mature pods representative of normal development were harvested from each plant grown under speed breeding II conditions. This resulted in 30 pods per genotype (3 replicate plants per genotype). Pods from each plant were placed together in 50 ml lidded plastic Falcon tubes with silica beads to absorb any moisture. These were then phenotyped using the pendulum pod shatter protocol [15] at the NSW Department of Primary Industries, Wagga Wagga Agricultural Institute (WWAI), Wagga Wagga, Australia. The rupture energy (RE) measures were adjusted for variation in pod length to calculate RELSQ (rupture energy per length squared), as per [25]. The five canola cultivars were also grown in field row plots at the experimental farm of WWAI in two replicates [16], and evaluated for shattering resistance using the same technique, where 10 fully mature pods were sampled per plot.

#### Wheat crossing and evaluation of crossing efficiency

Newly emerged spikes were emasculated to remove all immature anthers. The emasculated spikes were bagged for up to 3 days to allow for maturation of the stigma and then pollinated by anthers from a separately selected parent. The spikes were checked for seed setting 3-5 days post pollination. Crossing efficiency was calculated as the percentage of emasculated ovaries that successfully set seed.

#### Wheat flowering time

For measurement of flowering time of the wheat cultivars Paragon, Watkins landrace W352 [26], and the late flowering Paragon x W352 F_6_ recombinant inbred line, the number of days taken to reach heading was recorded for each replicate of each line. At the same time, the leaves on the main tiller were counted (starting from the 3-leaf stage) until the time of heading.

#### Wheat seed count

Seeds were manually counted in each spike. Images were obtained using a Nikon DSC D90 camera (HFX-DX, Melville, NY) and processed using ViewNX 2 software, produced by Nikon (Nikon, Surrey, UK) (Supplementary Fig. 12). Details are given in Supplementary Information.

#### Fusarium Head Blight infections

The test undertaken was for type II resistance, which is resistance to spread from floret to floret within the head. A suspension of 1x10^6^ conidiaspores per ml of *Fusarium graminearum*, which had been cultured in a mungbean liquid media, was used for point inoculations [27]. The isolate was a mix of two aggressive deoxynivalenol-producing *F. graminearum* UK isolates (K1-4/K1-5), obtained from the John Innes Centre’s Facultative Pathogen Collection. At least two spikes on each of three plants of the resistant line Sumai 3 and three plants of the susceptible line Timstein were inoculated.

#### Transformation of barley

Immature embryos of the barley *cv*. Golden Promise were inoculated with *Agrobacterium* strain AGL1 and transformed plants were recovered following the protocol of Harwood [28]. Constructs from the pBRACT series [29] containing selection cassettes conferring resistance to the antibiotic hygromycin were introduced.

#### Wheat meiosis studies

The anthers used for this study came from hexaploid wheat (*T. aestivum cv*. Chinese Spring (CS)) carrying the *Ph1* locus and crosses between rye (*Secale cereale cv*. Petkus) and hexaploid wheat (CS) either carrying or lacking the *Ph1* locus (*ph1b* deletion) [30].

Seeds were stratified on wet filter paper in the dark for 5 days at 4 °C, followed by a period of germination for 24 hours at 25 °C. The seedlings were vernalized for 3 weeks at 5 °C and then transferred to different controlled environment rooms until meiosis. Three seedlings from each genotype were transferred to the Conviron speed breeding chamber and an additional three were transferred to a glasshouse with no supplementary lighting or heating, as a control. From each plant, one spike was collected for the determination of meiotic stability in pollen-mother cells.

For meiosis studies, suitable young spikes were excised and fixed in ethanol:acetic acid (3:1). The samples were kept at room temperature, for at least one week, until slide preparation. Each floret has three synchronous anthers, and so one anther per floret was squashed in 45% acetic acid in water and assigned at metaphase I in meiosis by observation under a Leica DM2000 microscope (Leica Microsystems Ltd., Heerbrug, Germany; http://www.leicamicrosystems.com). The two remaining anthers were stained following the Feulgen procedure [31]. The present observations were limited to this stage. Images were collected using the Leica DM2000 microscope equipped with a Leica DFC450 camera (Leica Microsystems) and processed using LAS V4.4 system software (version 4.4, Leica Microsystems). Four parameters of meiotic stability were studied, including univalents, bivalents, multivalents and crossovers (COs). The number of cells observed in each plant in the two different photoperiods was 80. Further details are given in Supplementary Information.

Statistical analyses were performed using STATISTIX 10.0 software (Analytical Software, Tallahassee, FL, USA). The analysis of variance (ANOVA) was based on randomised blocks. In wheat-rye hybrids lacking the *Ph1* locus, tangent transformation was applied to univalents to meet the requirement of homogeneity of variances. Means were separated using the Least Significant Difference (LSD) test with a probability level of 0.05.

##### Acknowledgements

The authors wish to acknowledge the support of the Two Blades Foundation, USA and the Biotechnology and Biological Sciences Research Council (UK) grant numbers BB/H019820/1, BB/L009293/1, BB/L011794/1, BB/P013511/1 and BB/P016855/1, and the International Wheat Yield Partnership (grant IWYP76). SG was supported by a Monsanto Beachell-Borlaug International Scholarship. AW was supported by an Australian Post-graduate Award. MAMH was supported by a fellowship from Universiti Putra Malaysia, Malaysia. The authors also give thanks to the Australian Research Council for an Early Career Discovery Research Award DE170101296 to LTH and DAN00208 to HR. We are grateful to the JIC, UQ and PBI horticultural staff for plant husbandry, Michael Qiu (WWAI) for pod anatomy photography, Andrew Davis (JIC) for photography, and James Brown (JIC) for helpful discussions.

